# ProteinSGM: Score-based generative modeling for *de novo* protein design

**DOI:** 10.1101/2022.07.13.499967

**Authors:** Jin Sub Lee, Jisun Kim, Philip M. Kim

**Author notes:** Correspondence: Philip M. Kim.

## Abstract

The generation of *de novo* protein structures with predefined function and properties remains a challenging problem in protein design. Diffusion models, a novel state-of-the-art class of generative models, have recently shown astounding empirical performance in image synthesis. Here we use image-based representations of protein structure to develop ProteinSGM, a score-based diffusion model that produces realistic *de novo* proteins and can inpaint plausible backbones and domains into structures of predefined length. With *unconditional generation*, we show that ProteinSGM can generate native-like protein structures, surpassing the performance of previously reported generative models. We experimentally validate some *de novo* designs and observe strong structural consistency with generated backbones. Finally, we apply *conditional generation* to *de novo* protein design by formulating it as an image inpainting problem, allowing precise and modular design of protein structure.

## Introduction

Deep learning has significantly advanced protein engineering with robust methods for structure prediction and sequence design. AlphaFold 2 [1] allows researchers to access astronomically more protein structures while circumventing the strenuous effort of 3D structure determination, and recent sequence design methods [2, 3, 4] show robust sequence recovery given protein backbone information. One fundamental challenge in protein design that is largely unaddressed is the design of novel backbones - that is, first, can we generate synthetic backbones that can be realized by a protein sequence, and second, can we discover novel folds not found in the native fold space (i.e. folds that are not found in SCOP [5] or CATH [6])? An extension of this problem is the task of *conditional* backbone generation: for a given functional site, can we generate novel yet compatible scaffolds for the functional site that will retain its activity [7]? In this work, we seek to address these questions by leveraging recent advances in deep generative modeling.

Diffusion models [8, 9] have shown unprecedented success in various domains such as image synthesis [10], audio [11], text-to-image generation [12], graphs [13, 14], and small molecule generation [15]. Diffusion models define a forward diffusion process that perturbs data into noise and learns the reverse process that denoises random Gaussian noise into samples from the data distribution. Score-based generative modeling [9, 16] is a formulation of diffusion models that estimates the scores, or gradient of the log probability density with respect to data, of perturbed data in varying noise scales using a noise-conditional neural network. Sampling is performed by using the estimated scores with Langevin dynamics, which maps the Gaussian prior to the data distribution and thereby generates data from noise. This new generative model class has attained great empirical success due to its high sample quality, broad mode coverage, and increased training stability over other generative model types, and the use of diffusion models for protein design is now gaining significant attention [17, 18, 19, 20, 21]. Given these advantages and the proven empirical performance of diffusion models in various domains, we ask whether an appropriately engineered diffusion model trained on protein structural information can address various problems in protein design.

Here we present ProteinSGM, a continuous-time score-based generative model that generates high-quality *de novo* proteins. ProteinSGM learns to generate four matrices that fully describes a protein’s backbone, which are used as smoothed harmonic constraints in the Rosetta minimization protocol. Sequence and rotamer design is performed using fixed-backbone `FastDesign`, and lastly the structure is relaxed with constraints to generate a low-energy full-atom structure. We show that ProteinSGM generates variable-length structures with a mean < −3.9 REU per residue, indicative of native-like structures. We also provide an alternative approach that uses `MinMover` for backbone minimization, and ProteinMPNN [4] and OmegaFold [22] for sequence design and structure prediction. We experimentally validate the structural composition of a few generations with CD spectroscopy and find structural consistency between the generated models and experimental data. Moreover, we show that a conditional inpainting model is able to effectively inpaint masked coordinates, allowing direct control over protein functional sites while still generating realistic structures that follow the user-defined constraints.

## Results

In this work, we train a score-based generative model using image-like representations of protein structures, where each protein backbone is represented by inter-residue 6D coordinates [Figure 1A] as defined in trRosetta [23]. In brief, from each protein we calculate four matrices corresponding to C*β*-C*β* distances (hereafter referred to as *d*), *ω* and *θ* torsional angles, and *φ* planar angles that fully describe a protein backbone. These matrices constitute the 6D coordinates since *φ* and *θ* angles are asymmetric (ex. *φ_ij_* ≠ *φ_ji_* for residues *i, j* where *i* ≠ *j*). To generate proteins of different length, we also add a padding channel that indicates the boundaries of the 6D coordinates, which can be conditioned on by the model to generate proteins of fixed lengths. Taken altogether, a single protein structure is represented as a 128×128×5 tensor, with 4 channels corresponding to the 6D coordinates and one padding channel [Figure 1B].

**Figure 1.**
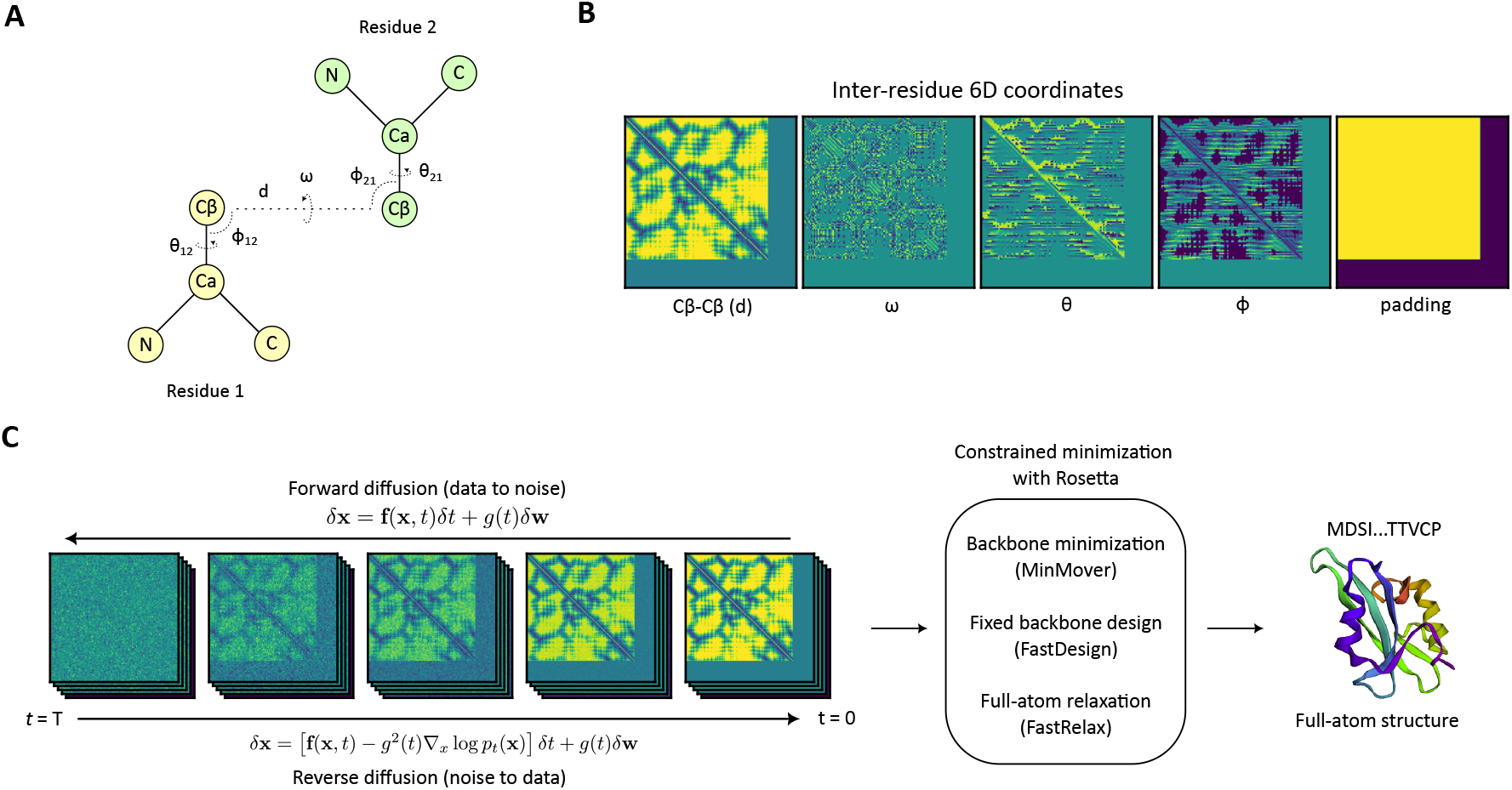
Model overview. (A) Inter-residue 6D coordinates *d, ω, θ, φ* between two residues. (B) Input features (6D coordinates and padding channel) used to describe a given protein structure. (C) A diffusion model is trained to generate realistic samples from noise by learning a reverse “denoising” process given the forward diffusion process that perturbs data to noise. The generated 6D coordinates are used as input for constrained minimization with Rosetta, which performed fixed-backbone design and full-atom relaxation to yield a protein structure corresponding to the 6D coordinate constraints.

We use the continuous-time framework of score-based generative modeling with stochastic differential equations (SDEs) [16], the first application to protein design to date [Figure 1C]. The model is trained to denoise realistic 6D coordinates from the Gaussian prior by estimation of the score function ∇_x_ log *p_t_* (*x*), which is used to solve the reverse-time SDE for mapping Gaussian noise into data (see *Methods*). The generated 6D coordinates are then subject to Rosetta minimization using `MinMover` for backbone minimization with constraints, `FastDesign` for fixed backbone sequence and rotamer design, and a final `FastRelax` constrained relaxation step to generate low-energy full-atom structures. After model training, we assess the model performance on unconditional generation to assess sample diversity and plausibility, and apply conditional generation by imputing masked input features for various protein design cases.

### Unconditional generation

#### 6D coordinate analysis

Adjacent residues of proteins constrain the inter-residue internal coordinates and therefore exhibit specific inter-residue distributions. To verify that the model is learning natural biophysical constraints of proteins and effectively capturing these distributions, we generated 1068 samples (12 samples for every length between 40-128) with the fully-trained model and compared 6D coordinate distributions of adjacent residues to the distributions of the test data [Figure 2]. Across all *d, ω, θ, φ* distributions, we observe that the distributions match closely to those of the test set, suggesting that the model has learned to generate realistic and native-like 6D coordinates of varying lengths. We also analyze joint distributions between 6D coordinates and observe that the adjacent residue features of true and generated samples are consistent across all 2D distributions [Supplementary Figure 1].

**Figure 2.**
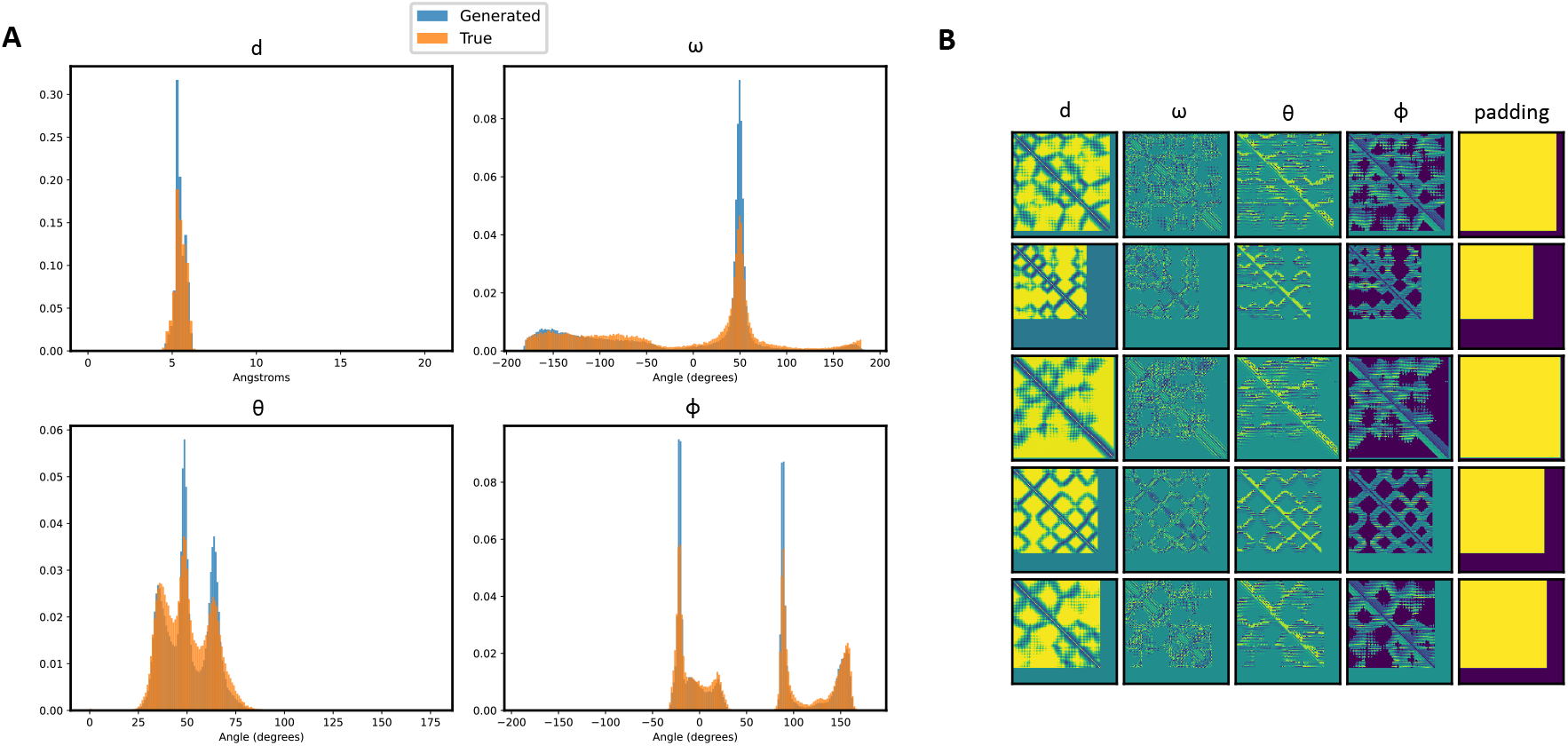
6D coordinate analysis. (A) 1068 samples were generated with the model and compared to features in the training set. *d, ω, θ*, and *φ* distributions of true (blue) vs generated (orange) samples show significant overlap, suggesting that the model has learned native-like constraints of interresidue 6D coordinates. (B) A few examples of the generated matrices.

#### Structure analysis

First, to verify the robustness of the Rosetta protocol in reproducible generation of structures from 6D coordinates, we extracted 6D coordinates from 750 structures in the test set and analyzed C*α* RMSDs on every minimization step [Supplementary Figure 2]. We observed that after `FastRelax`, the mean C*α* RMSD is 0.97Å, which suggests that the protocol is well-suited for reproducible generation at <1Å accuracy.

We proceeded to generate full-atom structures of all 1068 samples with the Rosetta protocol and compared their properties to minimized structures from the test set. We observed that the generated structures have Rosetta energies almost identical to those of native structures both with means of −3.9 REU per residue [Figure 3A], which is significantly higher than a recent VAE-based approach that reports a mean absolute REU of 0.061 with the best performing architecture [24]. To the best of our knowledge, this is the first generative model reported to show negative Rosetta energies across all generated samples. To detect generalization and diversity of the samples, we measured pairwise TM-scores with TMalign [25] between each generated sample and structures from the training set [Figure 3B]. We observed that a subset of the structures exhibited TM-scores of < 0.5, indicative of the model learning to generate novel folds not present in the training set [26]. Moreover, this indicates that ProteinSGM does not simply memorize structures found in the training set, which would strongly skew the max TM-score distribution to 1.0. When assessing the relationship between TM-score and Rosetta energy, we observed a strong negative correlation (*R*^2^ = −0.67), which indicates that generated structures with similar folds to native structures generally resulted in lower Rosetta energy, and therefore increased structural plausibility [Figure 3C]. As an additional validation of structural consistency across Rosetta minimization steps, we analyze the mean absolute error of the input 6D coordinates and 6D coordinates recovered after `MinMover` and `FastRelax`, and observe that d is conserved to sub-Angstrom accuracy, while the angular coordinates tend to vary up to 23 degrees after `FastRelax` [Table 1]. This may be explained by the increased flexibility of angular coordinates (especially torsion angles) compared to inter-atom distances, sub-optimal minimization by Rosetta, or inaccurate 6D coordinates that must be fixed during minimization. We observed that the generated 6D coordinates generally do exhibit greater deviations than structures minimized from native 6D coordinates, suggesting that Rosetta minimization indeed fixes some of the errors encountered during the diffusion process, which undoubtedly helps ProteinSGM generate structures of high fidelity. Therefore, Rosetta minimization can be viewed as a refinement step that optimizes the structure to higher quality. However, we note that after `FastRelax`, both native and generated 6D coordinates exhibit similar trends across all coordinates - for instance, *ω* displays a mean 23.2° and 18.8° deviation for generated and true samples, respectively.

**Figure 3.**
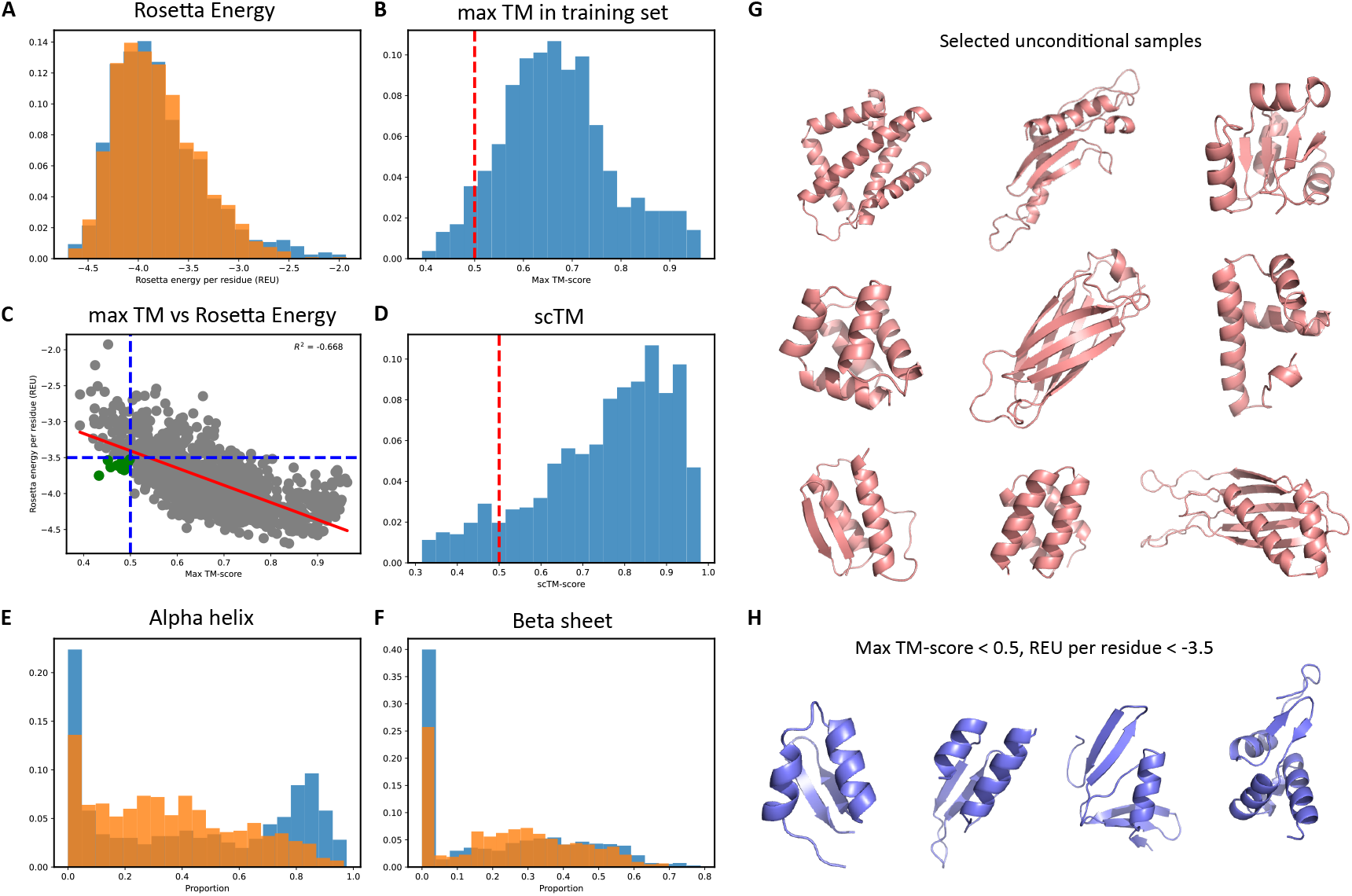
Structural analysis. 1068 generated samples were minimized, designed, and relaxed with Rosetta to produce full-atom structures. (A) Rosetta energies of the structures when compared to relaxed native structures show significant overlap with all generations exhibiting negative Rosetta energies, indicative of structural viability. (B) Max TM-scores were measured for each generation compared to all structures in the training set as a quantitative measure of generalization to novel folds not found in the training set. (C) There is a negative correlation between max TM-score and Rosetta energy, suggesting that structures with similar folds to the training set are generally more structurally viable, though a subset of structures (green) display max TM <0.5 and Rosetta energy per residue < 3.5, indicative of structurally plausible and novel samples. (D) An orthogonal assay with the ProteinMPNN sequence design model, OmegaFold, and the scTM metric shows that 90.2% of the structures (scTM > 0.5) can be realized by a protein sequence. Proportions of (E) *α*-helices and (F) *β*-sheets were measured between the generations and samples from the training set. The model tends to generate more helices and less *β*-sheets than native structures, which is expected since *β*-sheets are more difficult to generate due to long-range structural constraints. (G) A few examples of generated structures, including (H) ones that have novel folds (max TM-score < 0.5) and viable (Rosetta energy per residue < −3.5). More generations can be found in Supplementary Figure 5.

**Table 1.** Mean absolute error of input 6D coordinates and 6D coordinates recovered after `MinMover` and `FastRelax`.

|  | 6D coordinate | MinMover | FastRelax |
| --- | --- | --- | --- |
| Generated | $d$ ($C\beta - C\beta$) | 0.373Å | 0.469Å |
| | $\omega$ | 14.6° | 23.2° |
| | $\theta$ | 9.38° | 15.3° |
| | $\phi$ | 4.43° | 7.29° |
| True | $d$ ($C\beta - C\beta$) | 0.147Å | 0.349Å |
| | $\omega$ | 4.97° | 18.83° |
| | $\theta$ | 4.04° | 12.33° |
| | $\phi$ | 2.05° | 6.07° |

**Table 2.** Estimated secondary structure content (%) of unconditional samples by BeStSel [29]. Structure ID corresponds to the subfigure denoted in Figure 4.

|  |  | Structure ID |  |  |  |  |
| --- | --- | --- | --- | --- | --- | --- |
| Secondary structure |  | A | B | C | D | E |
| α-helix | Helix1 (regular) | 77.4 | 78.3 | 43.7 | 73.5 | 49.1 |
|  | Helix2 (distorted) | 21.8 | 21.1 | 23.2 | 17.1 | 21.6 |
| Anti-parallel β-sheet | Anti1 (left-twisted) | 0 | 0 | 0 | 0 | 0 |
|  | Anti2 (relaxed) | 0 | 0 | 0.8 | 0 | 0 |
|  | Anti3 (right-twisted) | 0.9 | 0.6 | 28.7 | 2.5 | 0 |
| Parallel β-sheet | | 0 | 0 | 0 | 0 | 0 |
| Turn |  | 0 | 0 | 0 | 1.5 | 0 |
| Others* |  | 0 | 0 | 3.5 | 5.3 | 29.3 |

\* 3,10-helix, $\pi$ -helix, $\beta$ -bridge, bend, loop/irregular and invisible regions

As an orthogonal method to assess designability of structures, we design sequences from sampled backbones (with `MinMover` prior to `FastDesign`) with the ProteinMPNN [4] sequence design model, OmegaFold structure prediction network [22], and the self-consistency TM (scTM) metric from [18] [Figure 3D]. We observe that 90.2% of the generations are designable given by scTM > 0.5, where a sequence (and predicted structure) with the same fold as the starting backbone can be generated. This is significantly higher than 11.8% and 22.7% reported in [18] and [19], respectively and is also comparable to scTM measurements of native structures, which yields 92.5% of structures with scTM > 0.5. We also analyzed the relationship between scTM and pLDDT - a structure prediction network’s confidence metric - since we reasoned that a reported scTM > 0.5 for a given backbone is less meaningful if the model prediction is not confident in the predicted structure [Supplementary Figure 3A]. We observed that when we remove predicted structures with mean pLDDT cutoffs at 70 and 90, corresponding to “confident” and “very confident” predictions, scTM proportions decrease to 78.3% and 24.5% for the generated samples and 88.9% and 48.3% for native structures, respectively. We also evaluated the relationship between scTM and max TM-score, and observed that there is a positive correlation (*R*^2^ = 0.60) between scTM and max TM-score [Supplementary Figure 3B]. This suggests that structures with higher similarity to the training set are generally more designable, which is expected since PDB structures in the training set are all realizable with a protein sequence. To analyze any length-dependent biases in the generated samples, we plotted scTM, max TM-score, and Rosetta energy per length [Supplementary Figure 4]. Interestingly, we see that for short length proteins, max TM-score is lower and Rosetta energy is higher than longer structures, with scTM not showing significant length biases. Intuitively, this makes sense since TM-align aligns the structures in the training set to the generated backbone prior to TM-score calculation, and therefore finding CATH domains in the training set that accurately match the generated folds is less observed. One explanation for the lower Rosetta energy is that smaller proteins exhibit less inter-residue interactions that stabilizes structural folds, thereby resulting in higher energies according to the Rosetta energy function.

Next, we analyzed the secondary structure distributions using DSSP [27] between native and generated samples [Figure 3E, F]. We noticed that the generated samples, when compared to native structures, have a marginally lower mean proportion of *β*-sheets (0.22 vs 0.24) and higher mean proportion of *α*-helices (0.44 vs 0.35), respectively. This is expected since *β*-sheets require specific local and global structural constraints for proper beta-sheet formation, while *α*-helices are more dependent on local interactions between neighboring residues and therefore easier to model, especially due to the inductive bias of convolutional networks. We present a few generated structures in Figure 3G, and structures with TM-score < 0.5 and Rosetta energy per residue < −3.5 in Figure 3H, which represent high-fidelity structures with novel folds not found in the training set. We also present two failure modes of ProteinSGM in Supplementary Figure 6: the generation of irregularly long helices and inaccurate *β*-sheet formation. Due to the abundance of helices in native proteins, most generative models, including ProteinSGM, has increased proclivity to generate helices as they are especially easier to model and therefore generate. In the case of inaccurate *β*-sheets, it is unclear whether this is an artifact of inaccurate 6D coordinate information, the Rosetta minimization protocol, or pyMOL misannotation. We note that in most cases, we observed that the *β*-sheets were recovered when the input backbones are passed through ProteinMPNN and OmegaFold, suggesting that the generated 6D coordinates were not the issue.

To ascertain that ProteinSGM can generate viable backbones that are foldable with a protein sequence, we selected a few samples for experimental validation [Figure 4]. First, we filtered minimized backbones from `MinMover` with a max TM-score < 0.6 to identify backbones with low structural similarity to the training set. After backbone candidate selection, we used ProteinMPNN to design 1024 sequences per input backbone and OmegaFold to obtain a predicted structure, from which we calculate scTM and extract pLDDT confidence scores. We filter the sequences by pLDDT >70 and rank each sequence by an average of pLDDT and scTM, and select the top 2 sequences per input backbone for experimental validation. We identify five candidates, all with max TM-score < 0.6 and high RMSDs to the closest example in the training set - with the exception of one instance in Figure 4E, where the closest training example is a helix-loop structure that exhibits little resemblance to the starting backbone. Circular dichroism (CD) is a popular method that uses differentially absorbed left and right circularly polarized light to distinguish secondary structural components of a given protein [28]. We sought to validate the generated structures using CD spectroscopy to validate the secondary structure composition of the designed proteins. To accomplish this, in addition to visual analysis of CD spectral peaks, we predict the fold composition from CD spectra using BeStSel [29]. Note that we observed strong consistency in results across both designed protein sequences per candidate backbone, and therefore we only show one representative plot per structure for clarity.

**Figure 4.**
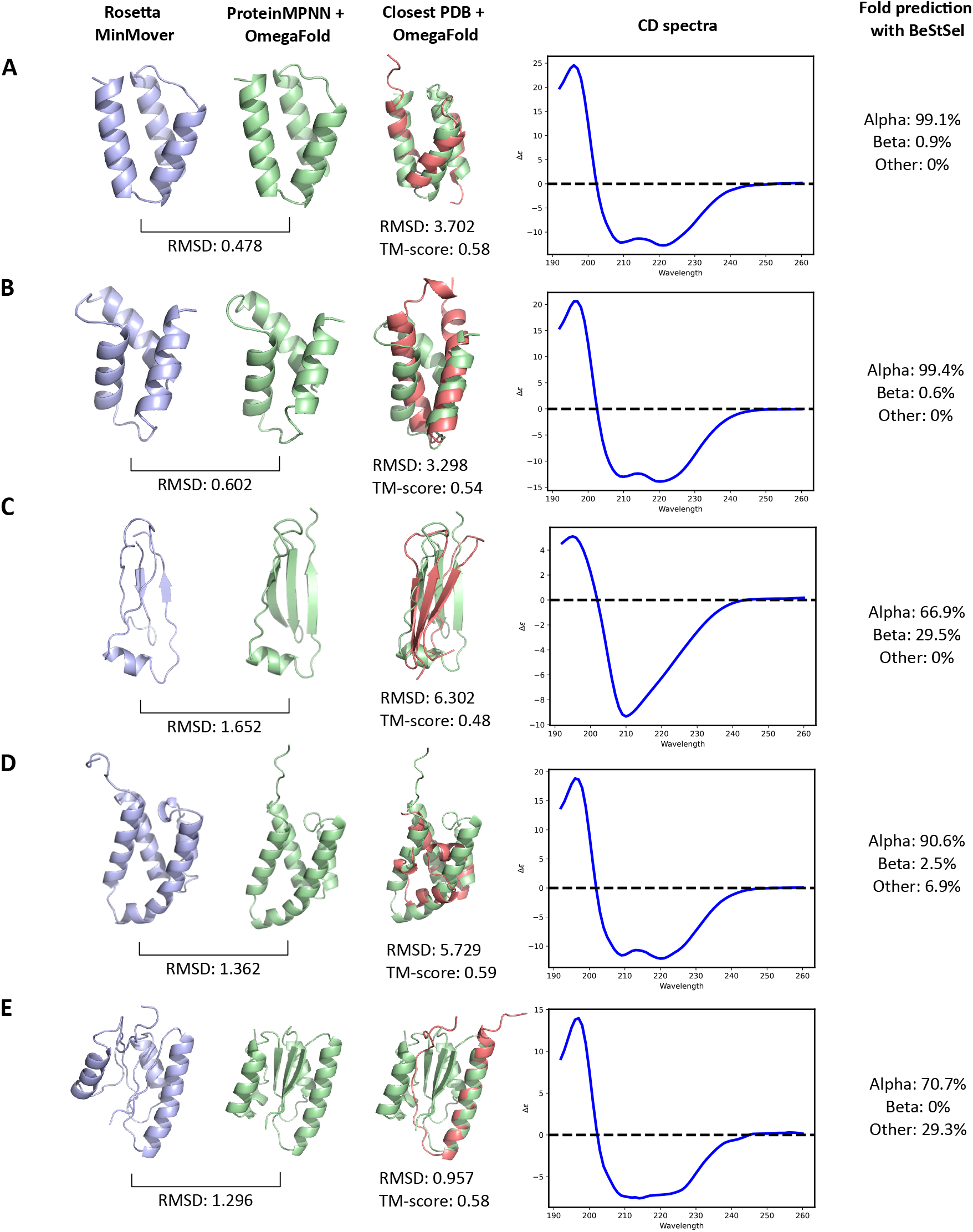
Experimental validation of unconditional samples generated by ProteinSGM, ProteinMPNN, and OmegaFold. We analyze five generations consisting of *α*, *α*β, and *β* proteins, and observe strong consistency between experimental CD spectra results and input backbone. We plot the experimental spectra and also analyze secondary structure predictions from CD spectra with BeStSel [29]. We observe that the *α* proteins (A,B,D) are generally very consistent, with positive peaks at 193 nm and negative peaks at 208 and 222 nm corresponding to typical *α*-only proteins. The *β*-dominant protein in (C) consisting of a novel fold with the max TM-score < 0.5 exhibits a strong negative peak at 210 nm that is characteristic of *β*-sheets. An *α*β-protein resembles a mainly *α*-protein, yet the absence of a bimodal peak at 208 and 222 nm and a stronger peak at 210 nm suggests the presence of *β*-sheets. Specific fold classification results from BeStSel can be found in Table 2.

In general, we observe that the *α*-helical structures [Figure 4A,B,D] show strong correspondence to expected values, where the experimental spectra closely resemble the patterns exhibited by typical *α*-only proteins with negative bands around 208 and 222 nm and a positive band around 193 nm [28]. We also analyzed one primarily *β* protein with a max TM-score < 0.5, and observed that the experimental spectra was more consistent with that of *β*-sheets with a strong negative peak at 210 nm. In the case of (E) that resembles a *α*-helical bundle surrounding *β*-sheets, we observe a strong positive band at 193 nm, negative bands at 208 and 222 nm indicative of *α*-helices, and a stronger negative signal at 210 nm corresponding to *β*-sheets, suggesting the presence of an *αβ*-protein. However, we note that BeStSel generally fails to predict *β*-sheets in fold prediction, which may be attributed to low proportion of beta sheets in this protein and the difficulty of generalizable deconvolution of CD spectra, especially given that the model was trained on a reference dataset of < 100 proteins that makes deconvolution of CD spectra corresponding to de novo folds especially difficult [29]. Overall, we find that structures generated by ProteinSGM are indeed designable, stably expressed, foldable, and show strong structural consistency through CD spectra analysis.

### Conditional generation

To address various tasks in protein design, we train a conditional diffusion model that learns to inpaint any masked region of a given input structure. The model is trained similarly to the unconditional model, except we introduce random masking of amino acids constituting 5-95% of the full protein. Specifically, we train the model in the unconditional setting (all residues masked), with random masking (random discontiguous masking), and with span masking (random contiguous masking), each sampled at equal probability at every training iteration (see *Methods*). This exposes the model to all potential conditional inpainting scenarios, and therefore can be applied to any protein design task.

Next, we present three practical protein design test cases for domain and scaffold inpainting. We used a recently published de novo-designed structure (PDB 2KL8) and masked out one helix domain of length 20 for input to the conditional diffusion model [Figure 5A]. We observed that across all generations, the model inpaints an *α*-helix to the masked region. This suggests that the model has learned, given the global structure constraints, that a helix can reasonably fit inside this pocket albeit with slight structural differences. This facilitates sampling structures with near-native topologies to optimize a functional property of interest, a central task in protein design. PDB 7MRX represents the bacterial barstar-barnase complex, an extensively-studied protein complex for its tight binding kinetics. Barstar inhibits the barnase ribonuclease primarily with a helix-loop domain for which we sought to design novel scaffolds [Figure 5B]. We identified regions to scaffold by masking residues whose C*α* distances were greater than 12Å from the target protein barnase. The generated scaffolds are diverse and show strong structural consistency between Rosetta structures and ProteinMPNN/OmegaFold structures at >80 mean pLDDT. This suggests that the generated scaffolds are structurally plausible for which sequences can be designed. In another test case, we sought to design scaffolds for the Mdm2, which inhibits the p53 tumor suppressor protein [Figure 5C]. Likewise we mask out residues whose C*α* distances were greater than 12Å from p53 and use the conditional ProteinSGM model to inpaint the masked regions. We generate strong candidates with high scTM and pLDDT that retain the binding site to p53 yet display diverse scaffolds, suggesting that the conditional model is suited for various scaffolding tasks. One potential application of scaffold inpainting for future exploration is the scaffolding of two disparate functional sites to generate synthetic bispecific proteins, which can be accomplished with ProteinSGM by imputation of scaffolds given two functional site descriptions.

**Figure 5.**
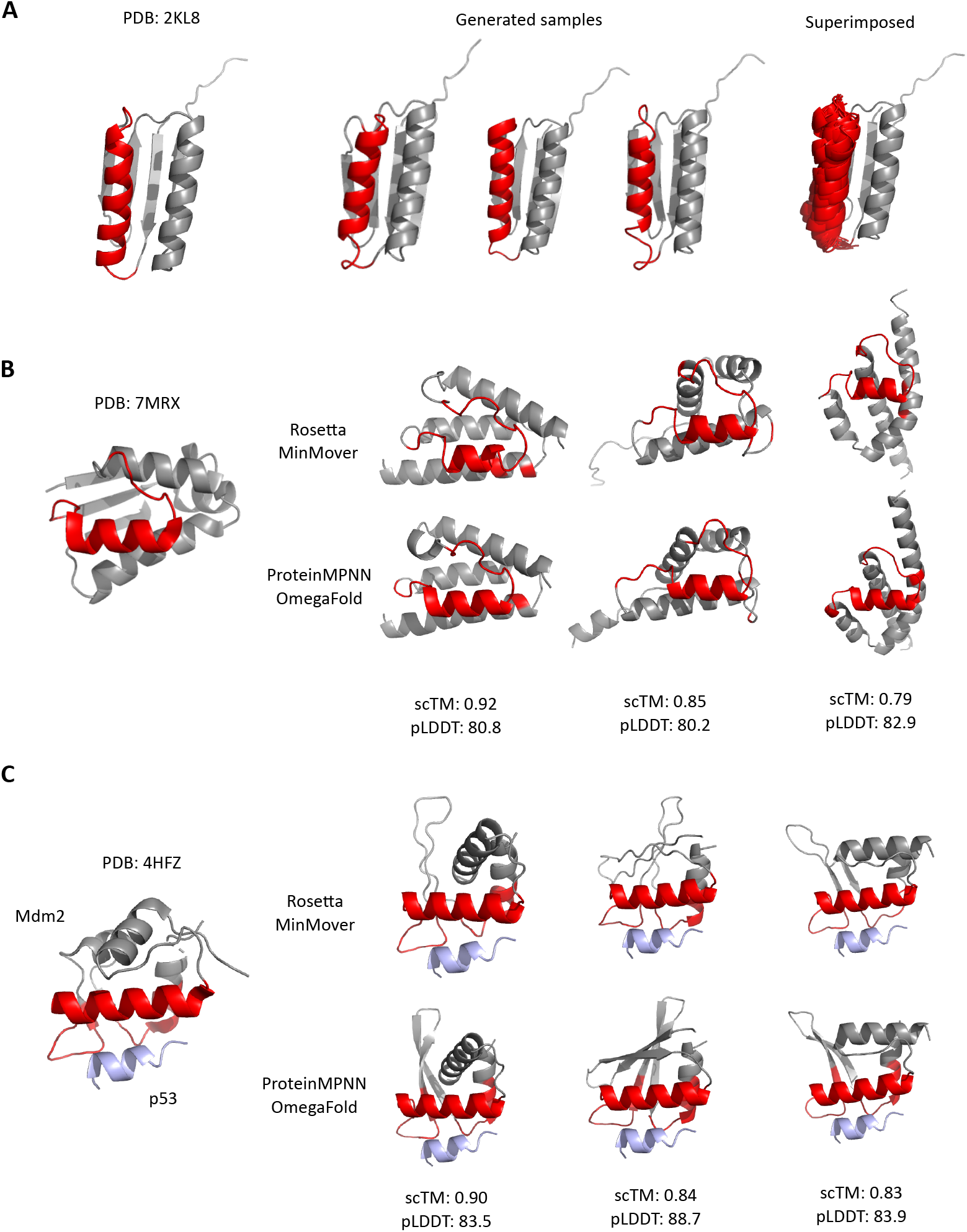
Protein design test cases. (A) As an example of domain inpainting, we ask the model to inpaint one helical domain of PDB 2KL8. Across all designs, the model inpaints slightly varied helices of similar shape to the original structure. The conditional inpainting model is tasked with inpainting the scaffolds of functional sites for barstar (B) and Mdm2 (C). We observe that the conditional model can generate viable scaffolds of varying topologies that retains the canonical barstar helix-loop domain and the native binding site of Mdm2 to p53. Rosetta structures and ProteinMPNN/OmegaFold structures show strong correspondence (scTM > 0.8 and mean pLDDT > 80), suggesting that these novel scaffolds are viable and realizable with a protein sequence.

We present an additional mode of conditional generation inspired by block adjacency conditioning from [17] [Figure 6]. We identify blocks as secondary structures identified by DSSP that correspond to *α*-helices or *β*-sheets in > 3 consecutive residues. We append two additional channels corresponding to the *α*-helix and *β*-sheet blocks, and a third block contact channel that indicates whether any C*α* coordinate between two blocks is within 7Å. This allows coarse conditioning of secondary structures, and therefore all generations conditioned on a given block adjacency description should consist of similar folds. Similar to [17], during training, we allow underspecification of the coarse block adjacencies by randomly dropping out blocks at a rate of 0.2. This allows further diversification of the generated structures without overconstraining the samples to a single fold. We present three coarse adjacency conditioning cases, and observe that across all cases, we can sample structures with increased diversity that remain structurally close to the native topology.

**Figure 6.**
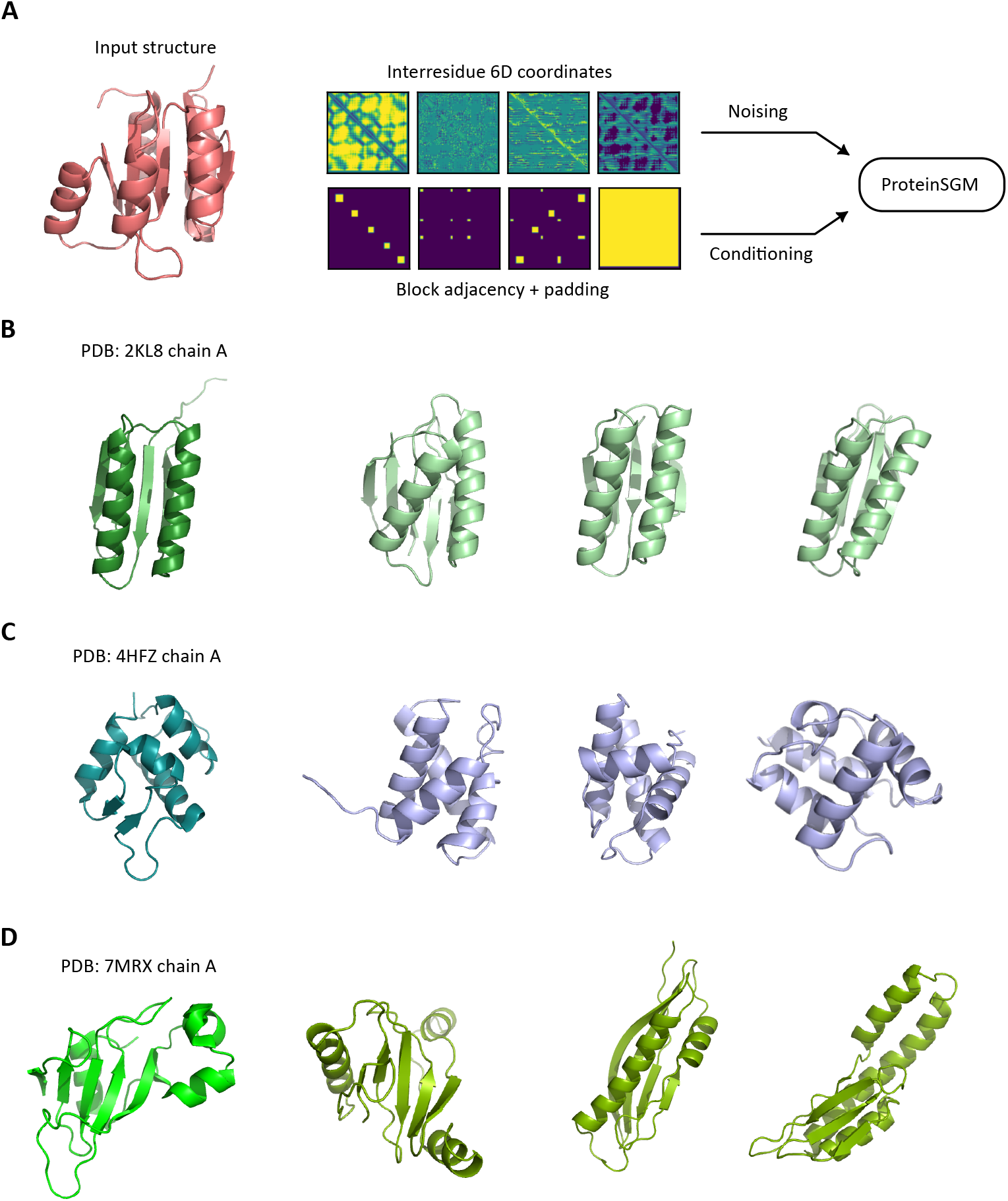
Block adjacency conditioning. (A) We train a block adjacency-conditioned model that learns to denoise 6D coordinates conditioned on coarse secondary structure descriptions. (B) When conditioned on the coarse adjacencies of 2KL8, we observe designs that look structurally close to the reference structure, yet are distinct - for instance, the left-most structure contains five antiparallel *β*-sheets, which is a distinct topology compared to the four found in 2KL8. (C) When conditioned on a helical structure, the model generates various helical structures that are uniquely distinguishable given the arrangement of the helices and the connecting loops. (D) In another test case of an *αβ*-protein, the generated samples are once again structurally diverse - note that the generations here are less constrained since the secondary structures are less packed, and therefore exhibit less contact constraints in the block adjacency channels.

## Discussion

This work presents one of the first applications of diffusion models to protein design - and the first to use the continuous-time SDE framework - that generates viable structures and can inpaint realistic backbones. The model learns to generate realistic 6D coordinates for which a structure is minimized with Rosetta. We obtain structures of native-like Rosetta energies that indicate structural plausibility, and observe that the generated structures are novel and diverse by TM-score analysis. We experimentally validate some generations by ProteinSGM and show structural consistency between generated structure and CD spectroscopy analysis. We apply the model for conditional generation, showing that the model can inpaint realistic domains and present novel scaffolds of varying lengths given functional site information.

Despite its high sample quality, one shortcoming of this approach is its computational cost, since sampling with the continuous-time diffusion model require many forward passes through the score network for solving the reverse SDE, and Rosetta relies on expensive MCMC procedures to traverse the energy landscape and find a local minima corresponding to a low-energy structure. We provide a systematic runtime analysis of all methods across all lengths in Supplementary Figure 7. While initial backbone minimization with `MinMover` is rather fast at < 2 minutes per structure, rotamer design and full-atom relaxation with `FastDesign` and `FastRelax` is very time-consuming and can take up to three hours per trajectory. Meanwhile, ProteinMPNN/OmegaFold takes up to 30 seconds per structure (on a single NVIDIA V100). Therefore, starting from 6D coordinate sampling, the estimated runtime of a single structure of length 100 is ~150 minutes with the full Rosetta protocol and 2-3 minutes with the `MinMover`/ProteinMPNN/OmegaFold protocol, which is up to 75-fold difference in runtime.

Moreover, while we do provide results with `FastDesign` and `FastRelax` for Rosetta energy calculations and structural visualization, we use the ProteinMPNN/OmegaFold pipeline for experimental validation due to its lower computational burden and the increased robustness of deep learning-based protein modeling. We present two approaches to full-atom structural generation, as the Rosetta-based method can be easily adapted to existing Rosetta optimization protocols, while the `MinMover`/ProteinMPNN/OmegaFold approach is more high-throughput and makes use of recent advances in deep learning for protein modeling. However, there are a few modifications to the Rosetta protocol that can be made to increase throughput. For instance, we suggest using a lower number of trajectories and/or `PackRotamersMover` for rotamer design, which significantly decreases compute cost with a marginal increase in Rosetta energy of the final structures. Though here we design the full-length protein from polyalanines, we note that for inpainting tasks where partial sequence and structure information are known, the Rosetta pipeline can easily be adapted to specifically design the inpainted region and retain the native domain(s). This significantly reduces runtime since only a subset of the structure is minimized and relaxed while keeping the rest of the structure fixed.

Wang *et al*. [7] reported the first case of using deep learning in scaffolding of functional sites by repurposing structure prediction networks (RosettaFold) to inpaint masked sequence and structural information. Moreover, several diffusion-based generative models for protein design have now been proposed, notably RFdiffusion [20] and Chroma [21]. Comprehensive comparisons of performance between these protein scaffolding and diffusion-based protein structure models are currently limited by availability of source codes as well as lack of a standardized dataset or metric to accurately measure and compare model performance like image synthesis (i.e. CIFAR dataset with FID metric). Ideally, experimental testing of novel designs should be conducted to properly assess design performance, but this is difficult to do at scale for comprehensive evaluation of different models. We also note that reconstruction of native proteins is often not an ideal metric as in inpainting approaches, as this only represent one possible solution of the design problem. Moreover, we emphasize that a key application of ProteinSGM is novel fold space exploration, since the unconditional ProteinSGM model is better suited for this task than recent models [18, 19], which exhibit low designability metrics and therefore is more prone to generate unrealizable proteins. Thus, these different approaches to protein structure modeling can be viewed as complementary yet distinct methods to address various protein design problems.

Recent developments have shown that diffusion models can be extended to large heteromeric protein complexes [21] and also for designing experimentally validated binders [20]. Likewise, potential improvements for ProteinSGM are to 1) increase modeling capacity to > 256 residues, and 2) incorporate multi-chain information for modeling protein-protein binding interactions, and 3) replacing Rosetta with a fully-differentiable module. Our initial efforts at increasing the maximum residue count to 256 were unsuccessful due to memory constraints, but with sufficient hyperparameter optimization and model tuning this should be attainable, especially since image-based diffusion models have shown strong empirical performance for resolutions up to 1024×1024 [30]. Incorporating multi-chain information would be beneficial for designing binders as shown in [20], allowing the model to learn implicitly learn binding interactions for effective conditional generation tasks such as antibody design. Finally, Rosetta minimization can be replaced with a structure generation module that takes in 6D coordinate information and generates the backbone coordinates to circumvent the time-consuming Rosetta protocol - we leave this for future work. One major limitation of current diffusion-based protein generation is that we still rely on the sequential generation of backbones, sequence, and rotamers, which restricts the natural flexibility of protein backbones and largely ignores sidechain-backbone interactions. Despite the exponential increase in computational complexity, extension of such generative models to one-shot full-atom generation is one promising avenue of work with great potential.

## Methods

### Dataset curation

We use the CATH [6] 4.3 40% sequence similarity clusters to reduce redundancy and potential bias in specific folds. After filtering for structures between length 40-128, we obtain 14,987 structures with 95/5 train/test splits. Note that we do not use experimentally-resolved C*β* coordinates, but rather extrapolate the position of the C*β* given the N, C*α*, and C coordinates. This also facilitates the modeling of glycines that do not have a *β* -carbon.

### Score-based generative modeling

For this work, we use the continuous-time framework of score-based generative models with stochastic differential equations (SDE). Here we provide a brief overview of the method - for more detailed information, please refer to Song *et al*. [16].

The forward noising process, given some data **x** and time *t* ∈ [0, 1], is defined by the following general-form SDE:

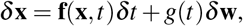

for **f**(·, *t*) is the drift coefficient, **g**(*t*) is the diffusion coefficient, and *δ***w** is white Gaussian noise. Intuitively, input data **x**, given a small time step *δt*, is noised by normally distributed random values with mean **f**(**x**, *t*)*δt* and variance **g**^2^(*t*)*δt*. Given a forward SDE, a corresponding reverse-time SDE can be formulated that requires knowledge of the score ∇_x_ log *p_t_* (**x**), or the gradient of the log probability density with respect to data, defined as follows:

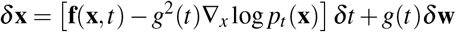

Therefore, once we define the forward noising process and learn an approximate score function, we can solve the reverse-time SDE and denoise random samples from a prior distribution into realistic samples. We use the VESDE (Variance Exploding SDE) discretization corresponding to the score matching objective with Langevin MCMC sampling and defined by the following SDE:

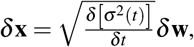

where σ(*t*) = σ_min_(σ_max_/σ_min_)*^t^* given lower and upper bound noise variance hyperparameters σ_min_ and σ_max_, respectively. To solve the reverse-time SDE, we estimate the score ∇*_x_* log *p_t_* (*x*) with a score network *s_θ_* (*x, t*). The score network is trained using a weighted denoising score matching objective [31], defined as follows:

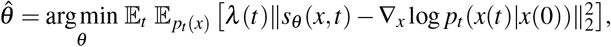

where *λ*(*t*) is a positive weighting function, and *p_t_*(*x*(*t*)|*x*(0)) corresponds to a perturbation kernel that perturbs clean sample *x*(0) to noisy sample *x*(*t*).

We retained the UNet-based architecture [32] for score estimation as used in [16] with multi-head self-attention in the 16×16 resolution block. We reasoned that using the UNet architecture for the score network is justified since the task is presented as an image synthesis problem, and the input data resembles image-like data with 5 channels. For the noise schedule hyperparameters, we heuristically use σ_min_ = 0.01 and σ_max_ = 100 after brief optimization. The model is trained with a single NVIDIA V100 GPU using a batch size of 8 and learning rate 1 × 10^-4^ for 2 million iterations, which consumes approximately 7 days.

Sampling is performed by using Predictor-Corrector sampling (Reverse diffusion Predictor with Langevin dynamics Corrector) which provides an initial prediction for a given denoising step using the estimated scores from the score network, and is further refined using Langevin dynamics. We use *N* = 2000 timesteps for sampling.

### Rosetta minimization

To obtain structures from the inter-residue 6D coordinates, we use an adaptation of the trRosetta minimization protocol. While trRosetta and a previously reported VAE-based approach [24] use distograms, or probabilities of each distance/angular bin, to fit a spline function to generate smoothed constraints for Rosetta minimization, ProteinSGM directly generates the distance and angular values. We then use the HARMONIC function for *d* and *φ* and CIRCULARHARMONIC for *ω* and *θ*, with the mean corresponding to the generated value and the standard deviation set to 2.0Å for d and 10° for *φ, ω*, and *θ*. This allows reproducible generation of the structure given a set of 6D coordinates while still relaxing the constraints enough to generate realistic structures [Figure S1].

Since we do not have sequence information for the generated matrices, we use the padding channel to obtain the matrix boundaries *L* by *L* of the generated matrices, and a polyalanine chain of length *L* for the minimization protocol. We also use the upper triangle for the *d* and *ω* matrices since they are symmetric, and do not include any constraints that are *d* > 12Å apart.

To generate backbone from constraints, we use `MinMover` with 5 rounds of minimization and select for the pose with lowest energy using a coarse-grained centroid energy function. Each round of minimization progressively uses short, medium, and long-range constraints by sequence separation, and randomly changes the *ω* and *φ* backbone torsional angles by up to 10 degrees. Once a tentative backbone has been generated, we use `FastDesign` for fixed-backbone design to sample different sequences and rotamers that can fit the generated backbone. Then we idealize problematic regions and perform full-atom relaxation of the structure using the matrix constraints. The entire process is repeated for 3 independent trajectories, and the lowest energy structure (using the `ref2015` energy function [33]) is selected as the final structure.

All Rosetta protocols were written with `pyRosetta` 4.0 [34].

### Sequence design and structure prediction

As an alternative method to survey designability of generated backbones, we adapt the self-consistency TM (scTM) metric from [18]. After backbone selection from `MinMover`, instead of passing the backbone to `FastDesign`, we use the ProteinMPNN [4] sequence design model with temperature = 0.1 to sample 8 sequences for each backbone, and predict structures for each sequence with OmegaFold [22]. We calculate the TM-score between each OmegaFold-predicted structure and the starting backbone, and the highest TM-score across all 10 structures is designated as the scTM score for the given backbone. Backbones with scTM > 0.5 are considered designable structures, since at least one sequence can be designed for which a predicted structure has the same fold (TM > 0.5) as the starting backbone.

### Conditional model training

We present two conditional models adapted from the unconditional ProteinSGM model. To create a general inpainting model, rather than noising and denoising all 6D features, during training we mask 5-95% of the residues with either random masking or span masking. In random masking, we sample a random probability *p* where 0.05 ≤ p ≤ 0.95, and every residue is masked with a probability p. Note that this is a soft masking, as in the actual proportion of masked residues may not equal *p*. In span masking, given a protein of length *L*, we sample a random integer *r* where 0.05*L* ≤ *r* ≤ 0.95*L*, sample a starting residue i where 0 ≤ *i* ≤ *L* – *r*, and mask positions [*i, i + r*]. On every training iteration, the masking strategy - unconditional, random masking, and span masking - is sampled at equal probabilities, so that the model is exposed to all cases of conditional generation during training.

For the block adjacency conditioned model, we extract the block adjacencies similar to [17], where each *α*-helix and *β*-sheet block is identified using DSSP, excluding blocks that are < 3 residues long. The contact channel is generated by iterating over all pairs of block adjacencies, and identifying a block contact if any pairwise C*α* distance between two blocks is < 7Å. Then at every training iteration, individual blocks are dropped out at a rate of 0.2 to allow underspecification of the coarse block adjacencies. We do not dropout any blocks during inference.

### Protein purification

The DNA fragments encoding designed proteins, were prepared by DNA synthesis (IDT technology). For mammalian expression, each DNA encoding designed proteins was cloned in frame into a pcDNA3.1 vector, which include a signal peptide and a N-terminal 8× His-tag. The plasmid was transiently transfected into cultured HEK293T cells in the serum-free DMEM medium following the standard protocol. Culture supernatants were collected after 3-6 days by centrifugation. Proteins were purified from the culture supernatants on a Ni-NTA Superflow (Qiagen) and were extensively dialyzed to achieve the final composition of PBS buffer and their concentrations were determined by measuring the absorbance at 280 nm.

### Circular dichroism (CD) analysis

The CD spectra (192-260 nm) of the unconditionally designed proteins (0.5 mg/mL in PBS buffer, pH 7.4) were analyzed on a J-810 circular dichroism spectropolarimeter (JASCO Inc.) at 25°C, in a 1 mm path length quartz cuvette with a step size of 1 nm. Averaged spectra of ten scans were corrected for buffer blank (PBS buffer, pH 7.4) and expressed as delta epsilon (Δ*ε*).

## Code and Data Availability

The codebase and dataset used for this work is available at https://gitlab.com/mjslee0921/proteinsgm.

## Author Contributions

J.S.L. and P.M.K. conceptualized the work, J.S.L. developed the computational results, J.S.K. performed the experimental validation and J.S.L., J.S.K., and P.M.K. wrote the manuscript. P.M.K. supervised the work and acquired funding.

---

**Figure S1.**
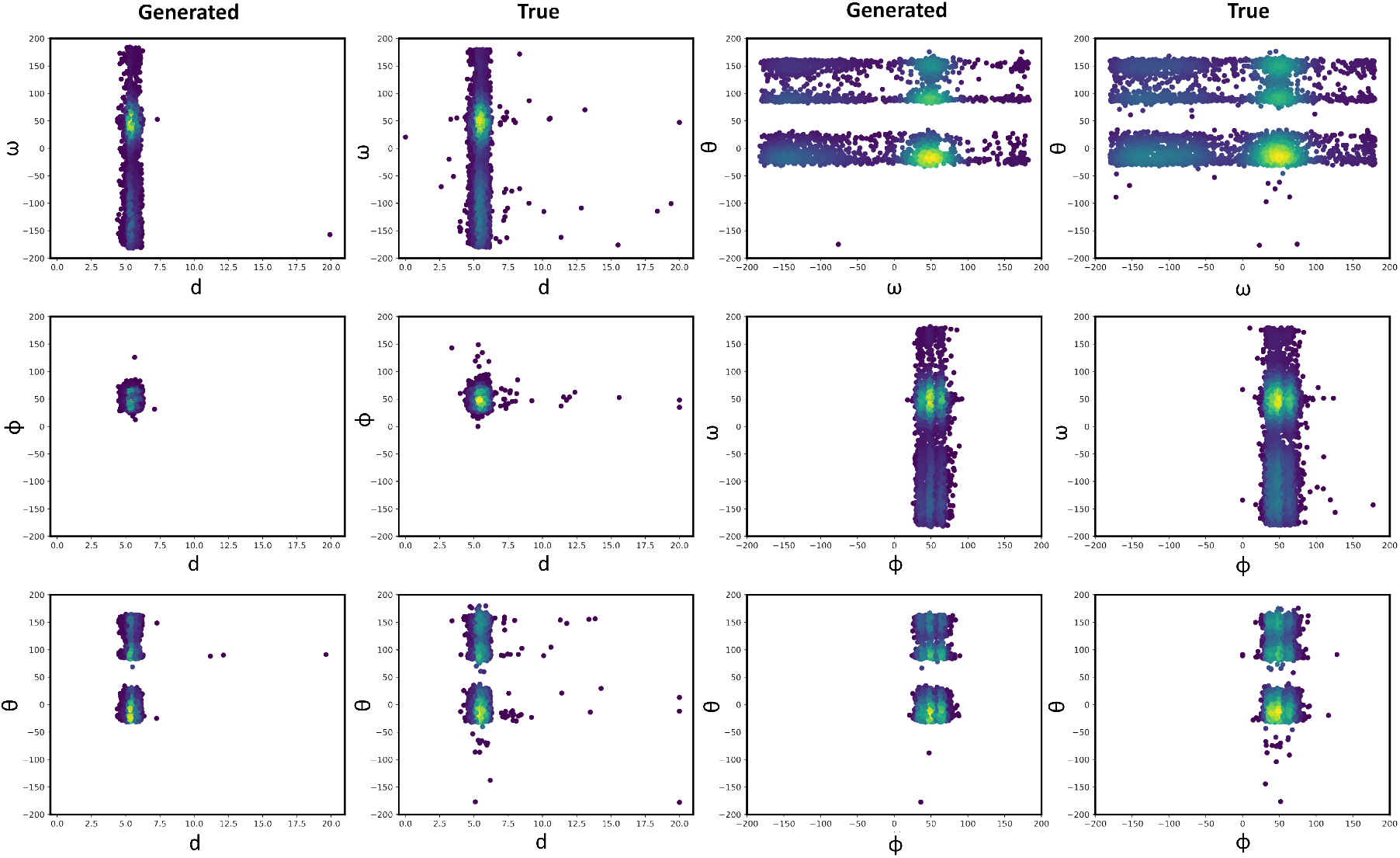
2D matrix distributions. We analyzed joint distributions of 6D coordinates in adjacent residues between true and generated samples to further assess generation quality of ProteinSGM. We see a clear concordance between generated and true samples, suggesting that ProteinSGM has learned native biophysical constraints.

**Figure S2.**
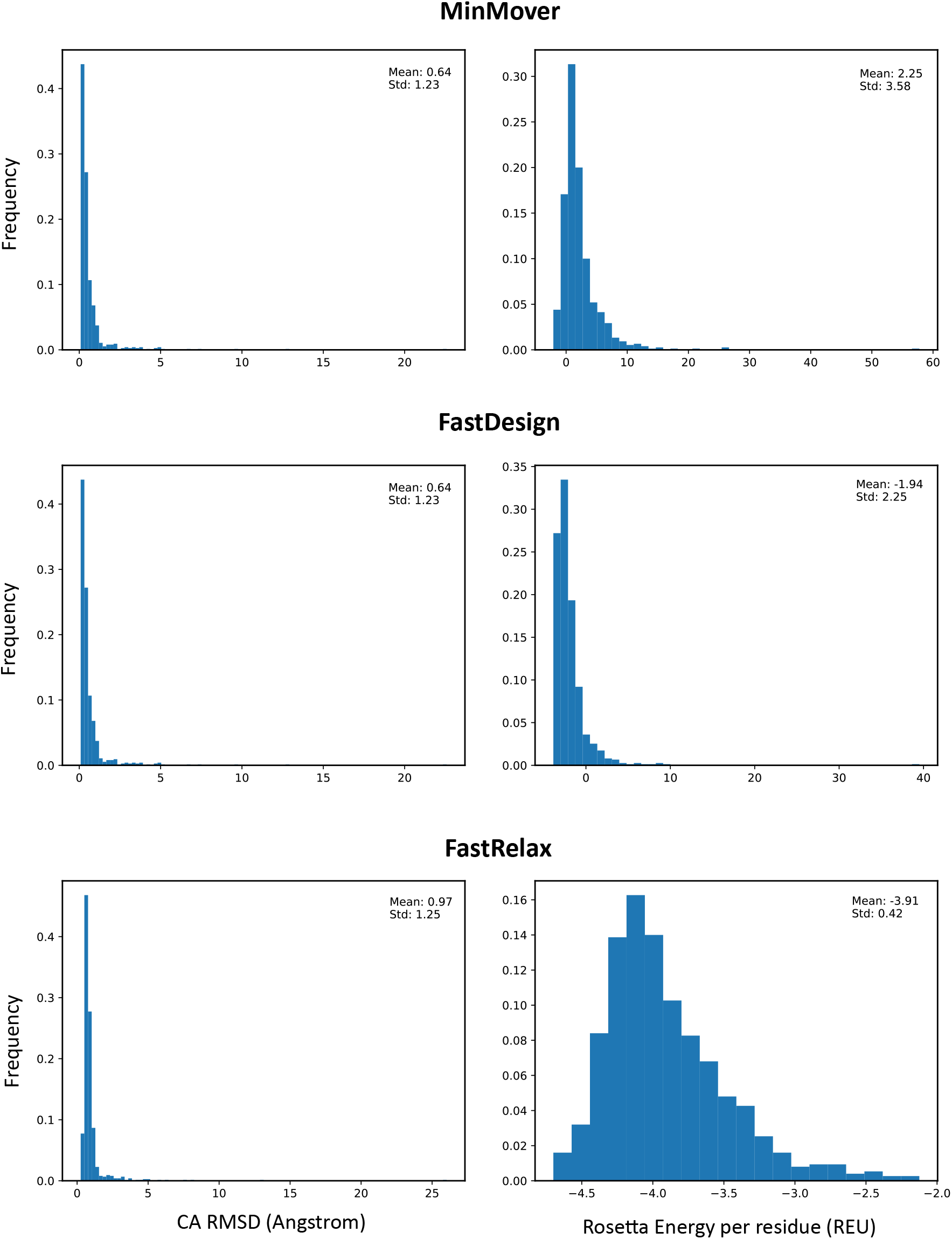
Rosetta energies and C*α* RMSDs with Rosetta pipeline. (left) Rosetta energy before `FastRelax` remains relatively high for generated structures, but full-atom relaxation with constraints allows effective energy minimization to bring energies closer to the true distribution. (right) Minimized structures from 6D coordinates were compared with their native structures to assess fidelity of the Rosetta protocol. We observe that across all steps, mean C*α* RMSDs are lower than 1Å, suggesting the protocol is suited for reproducible generation of structures from 6D coordinates.

**Figure S3.**
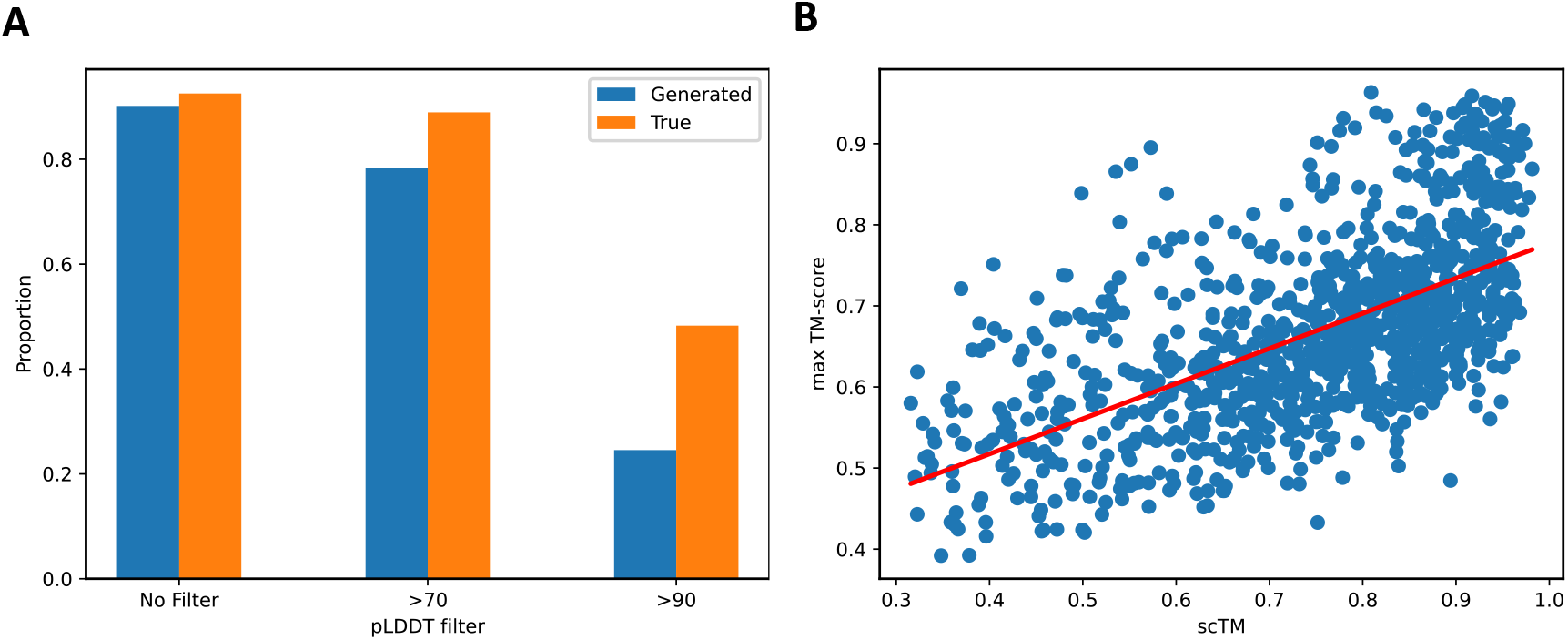
Additional scTM analysis. (A) Since predicted structures with low pLDDT (low confidence) are generally unreliable, we filter OmegaFold structures at different cutoffs and analyze changes to the scTM > 0.5 proportion. With no filtering, a > 70 pLDDT filter, and a > 90 pLDDT filter, we obtain 90.2%, 78.3%, and 24.5% scTM > 0.5 in the generated samples and 92.5%, 88.9%, and 48.3% in the true samples, respectively. Though there is a notable difference between generated and true samples with more stringent pLDDT filters, we observe that at a > 70 pLDDT filter corresponding to confident predictions, more than 78% of the generated samples are designable with a protein sequence. (B) We analyze the relationship between scTM (designability) and max-TM (similarity to training data), and observe that there is a positive correlation between scTM and max TM-score. This is expected since structures with closer similarity to PDB structures are by definition realizable with a protein sequence, and therefore should have an scTM > 0.5.

**Figure S4.**
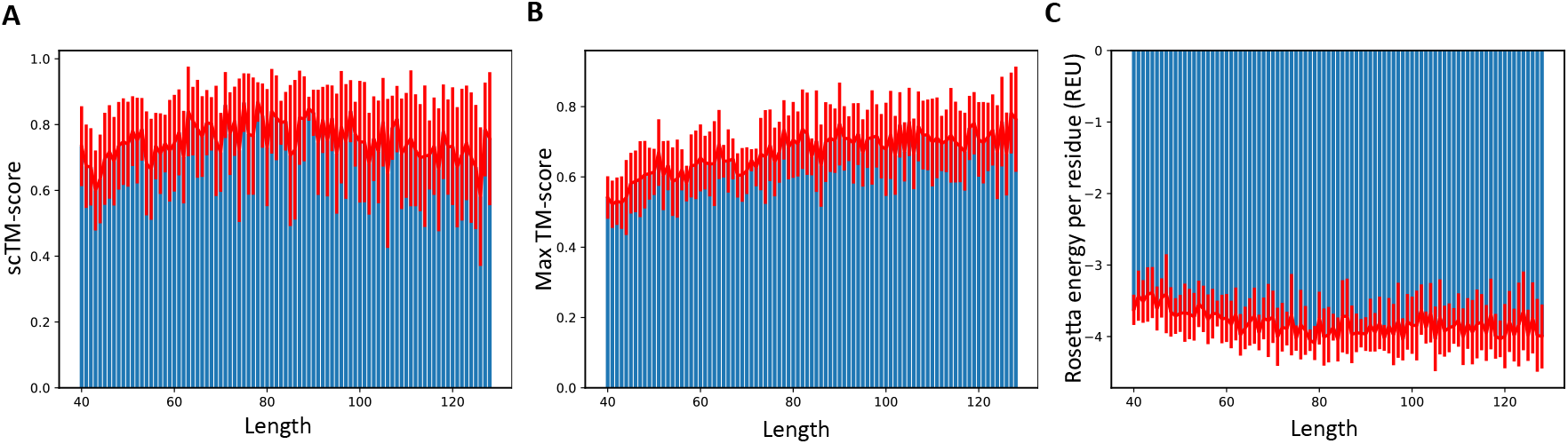
Per-length analysis of (A) scTM, (B) max TM-score, and (C) Rosetta energy. We observe that there is consistency across all metrics between structures of different lengths. However, there is a slightly lower max TM-score and higher Rosetta energy for shorter structures, which may be explained by the lower number of short domains in the training set and the lower number of stabilizing interactions in shorter proteins, respectively.

**Figure S5.**
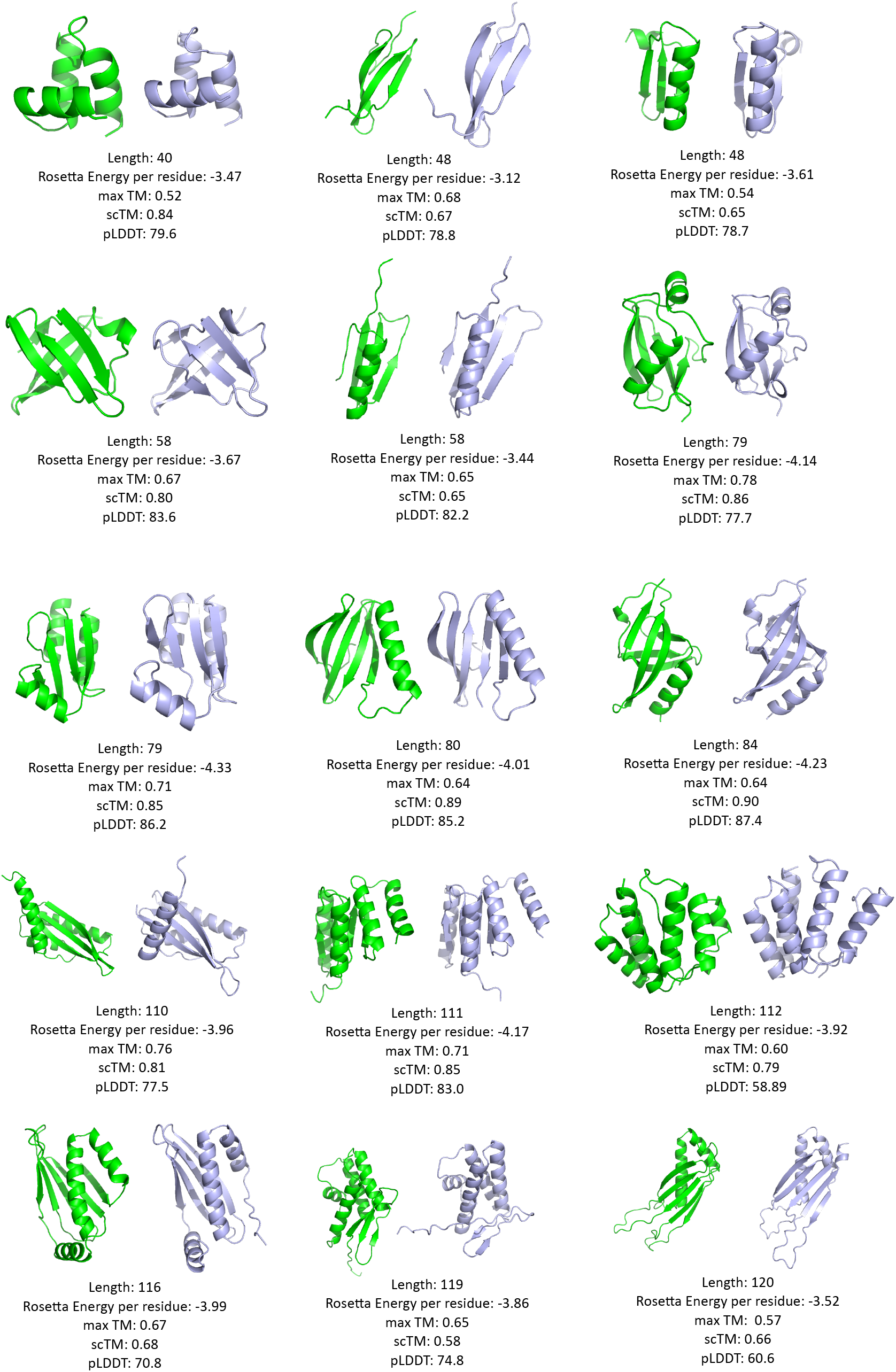
Examples of generated structures. Rosetta-generated backbones are in green, and the best OmegaFold structure is shown in blue.

**Figure S6.**
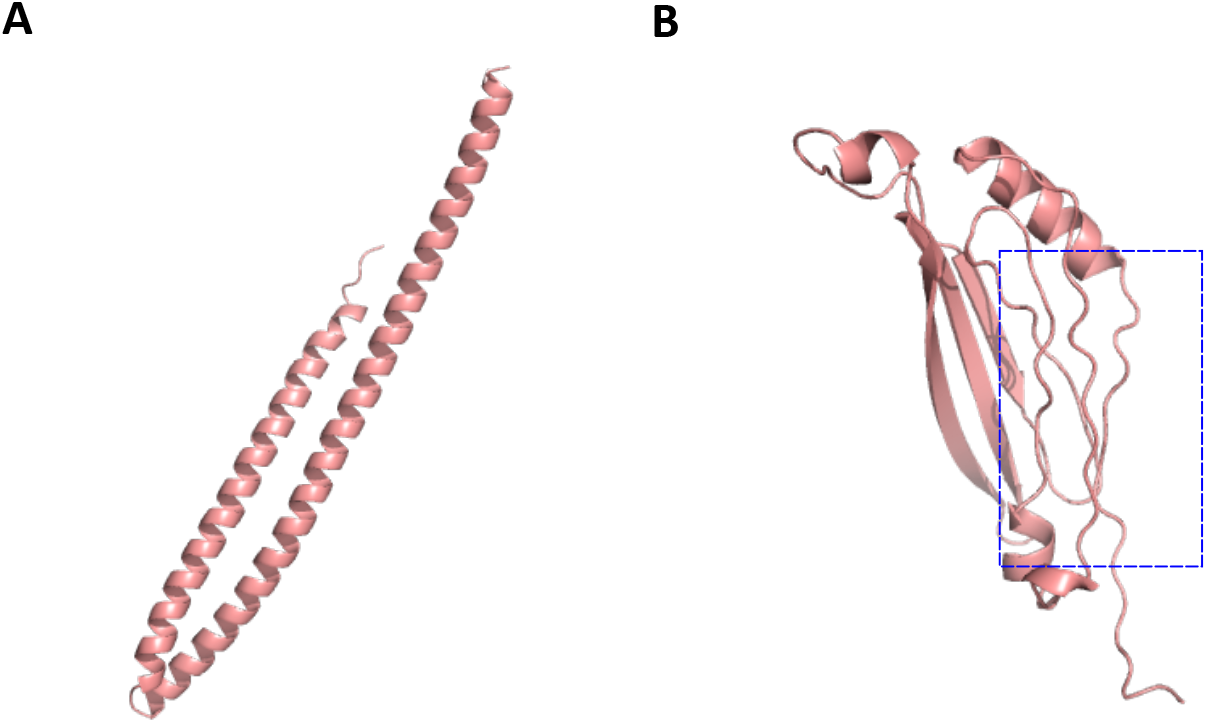
Failure modes of ProteinSGM. (A) We observe that in a few cases, ProteinSGM generates long helices that may not exist in nature. (B) Since *β*-sheets are more difficult to model, some generated backbones display loop-like structures that are not visualized as *β*-sheets due to incorrect placement of hydrogen bonds.

**Figure S7.**
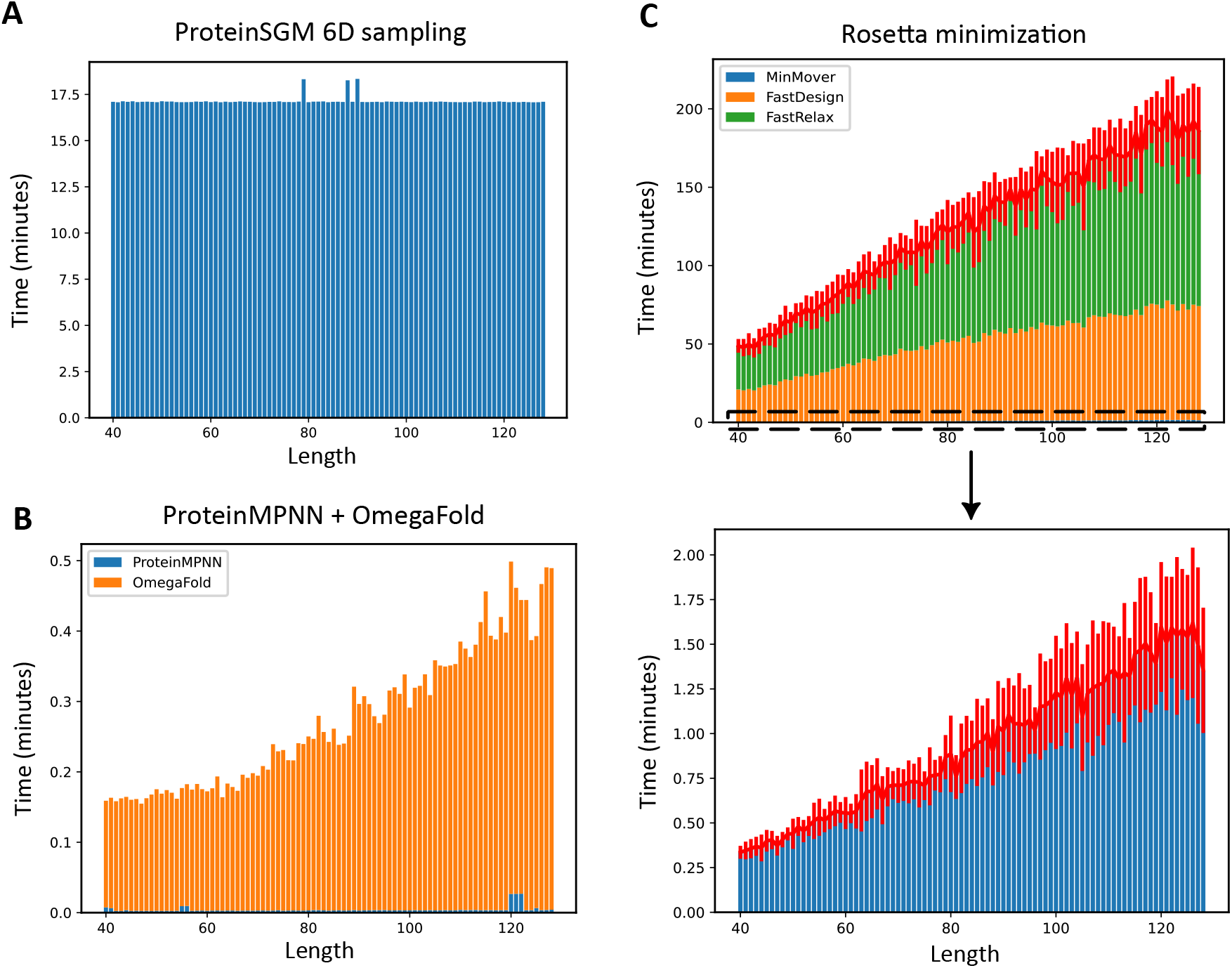
Runtime analysis by length. (A) Sampling from ProteinSGM is uniform across all lengths since the dimensionality of the generated matrices is fixed. (B) With the ProteinMPNN + OmegaFold pipeline, OmegaFold dominates most of the runtime, yet is relatively fast at < 30 seconds per structure. (C) With the Rosetta pipeline, initial backbone generation with `MinMover` is relatively fast at < 2 minutes per structure, but `FastDesign` and `FastRelax` can take up to 3 hours per structure. ProteinSGM, ProteinMPNN, and OmegaFold runtimes were measured using a single NVIDIA V100, while Rosetta was run with 2 cores and 8GB of RAM. Standard deviations for Rosetta iterations are shown in red. ProteinSGM was sampled with batch size 16, and ProteinMPNN was run in batch mode with 12 input structures and 8 sampled sequence per structure, so runtime may vary with different batch sizes.

